# Bone Morphogenetic Protein (BMP) signaling upregulates expression of ID1 and ID3 in pancreatitis and pancreatic ductal adenocarcinoma

**DOI:** 10.1101/2023.09.01.555987

**Authors:** Megha Raghunathan, Kathleen M. Scully, Amanda Wehrmaker, Rabi Murad, Andrew M. Lowy, H. Carlo Maurer, Kathleen E. DelGiorno, Pamela Itkin-Ansari

## Abstract

Identification of biological modulators of pancreatic ductal adenocarcinoma (PDAC) initiation and progression is of critical importance as it remains one of the deadliest cancers. We have previously shown that ID1 and ID3 are highly expressed in human PDAC and function to repress expression of E47 target genes involved in acinar cell differentiation and quiescence. Here, mining of large bulk RNA-seq and single cell RNA-seq (scRNA-seq) datasets has associated high expression of ID1 and ID3 with poor PDAC patient survival. We show that upregulated expression of ID1 and ID3 in human PDAC cells occurs in response to canonical BMP signaling through pSMAD1/5/9. Conversely, treatment with Noggin, an endogenous BMP antagonist, or with DMH1 and LDN193189, small molecule inhibitors of BMP receptors, reduced expression of ID1, ID3 and cell cycle control genes. Based on our RNA-seq and immunohistochemical analyses, upregulation of BMP signaling to ID1 and ID3 is an early event that occurs in human pancreatic intraepithelial neoplasia (PanIN) and murine models of pre-neoplastic lesions induced by mutant Kras. Strikingly, the same result was observed in a murine model of pancreatitis induced by the cholecystokinin (CCK) analog caerulein. Moreover, we show that caerulein is sufficient to induce BMP signaling and expression of ID1 and ID3 in a cell-autonomous manner in non-transformed rodent exocrine pancreas cells. Together, the data suggest that canonical BMP signaling upregulates expression of ID1 and ID3 early in pancreas pathogenesis and that pancreatic cancer cells remain addicted to this important signaling circuit as the disease progresses. Future exploration of druggable targets within this pathway could be of therapeutic benefit in the treatment of pancreatitis and PDAC.

## Introduction

ID1-4 (inhibitors of DNA-binding and differentiation) comprise a family of helix-loop-helix (HLH) proteins. Their primary function is to interact with basic helix-loop-helix (bHLH) transcription factors to prevent them from forming dimers that bind to target genes. ID proteins inhibit cell differentiation, promote DNA replication and cell survival, and can subvert KRAS-induced senescence ^1^. High levels of ID1 and/or ID3 are associated with a variety of cancers, including ovarian cancer, thyroid cancer, and glioblastoma ^2–4^.

We and others showed that human and murine pancreatic ductal adenocarcinoma (PDAC) cells expressed ID1 and ID3 and that knockdown reduced cell growth ^5–11^. We also demonstrated that ID3 overexpression was sufficient to induce cell cycle entry in normally quiescent human acinar cells, a cell of origin for PDAC^6^. Mechanistically, we found that ID1 and ID3 bound to the bHLH transcription factor E47 (a splice variant of E2A) which resulted in suppression of E47 target genes required for acinar cell differentiation and quiescence ^7^. Important questions remain about how expression of ID1 and ID3 are induced in PDAC. Moreover, because protein-protein interactions (e.g. between IDs and E47) are notoriously difficult to target pharmaceutically, it is critical to identify potentially druggable pathways that impinge on expression of ID1 and ID3 in PDAC.

Bone morphogenic proteins (BMPs) belong to the transforming growth factor-β (TGF-β) superfamily. This is a large group of structurally related signaling proteins including soluble BMP ligands. These ligands bind to specific BMP type I receptors ALK1 (ACVRL1), ALK2 (ACVR1), ALK3 (BMPRIA), or ALK6 (BMPRIB) that heterodimerize with and phosphorylate one of three BMP type II receptors (ActRII, ActRIIB or BMPR2). In canonical BMP signaling, the activated ALK1/2/3/6 receptors phosphorylate R-SMAD1/5/9 proteins that form a complex with SMAD4 to enter the nucleus and act as a sequence-specific DNA-binding complex to regulate target gene transcription.

In this study, we investigated the relationship between BMP signaling and expression of ID1 and ID3 in diseased pancreatic epithelial cell populations using data from human patients, human PDAC samples, mouse tissue, and human and rat cell lines. We also examined earlier stages of pancreas pathogenesis in preneoplastic pancreatic intraepithelial neoplasia (PanIN) lesions and in a murine model of chronic pancreatitis, itself a major risk factor for PDAC. Overall, the data revealed a highly conserved BMP-stimulated increase in ID1 and ID3 expression that begins early, is sustained throughout the course of pancreatic disease, and is correlated with poorer outcomes for patients.

## Results

### High expression of ID1 and ID3 in human PDAC is associated with shorter patient survival times

Previously, we and others showed upregulation of ID1 and ID3 protein in immunostained human PDAC tissue samples^5–8^. Analysis of numerous published large-scale bulk RNA profiling studies that compared hundreds of human PDAC tumors to normal pancreas or tumor adjacent pancreas tissue allowed us to ask focused questions about the expression of ID1 and ID3 mRNA in large numbers of patients. Comparison of ID1 and ID3 mRNA expression in five RNA-seq datasets of normal versus PDAC tissue revealed statistically significant increases in both genes in PDAC in all datasets examined (GSE16515, GSE28735, GSE60979, GES62452, CPTAC)^12–16^. The number of normal samples in each dataset ranged from 12-61 and the number of PDAC samples from 36-140. In all five datasets, the difference between the medians for normal and PDAC was greater for ID1 than ID3 (**Figure 1A**). In contrast to IDs, the level of E2A (*TCF3* gene) in the 5 datasets was not consistently up or downregulated in PDAC, supporting our previous findings that E2A is primarily dysregulated by ID proteins at the level of function, not expression **(Figure S1)** ^12–15,17^.

**Fig. 1:**
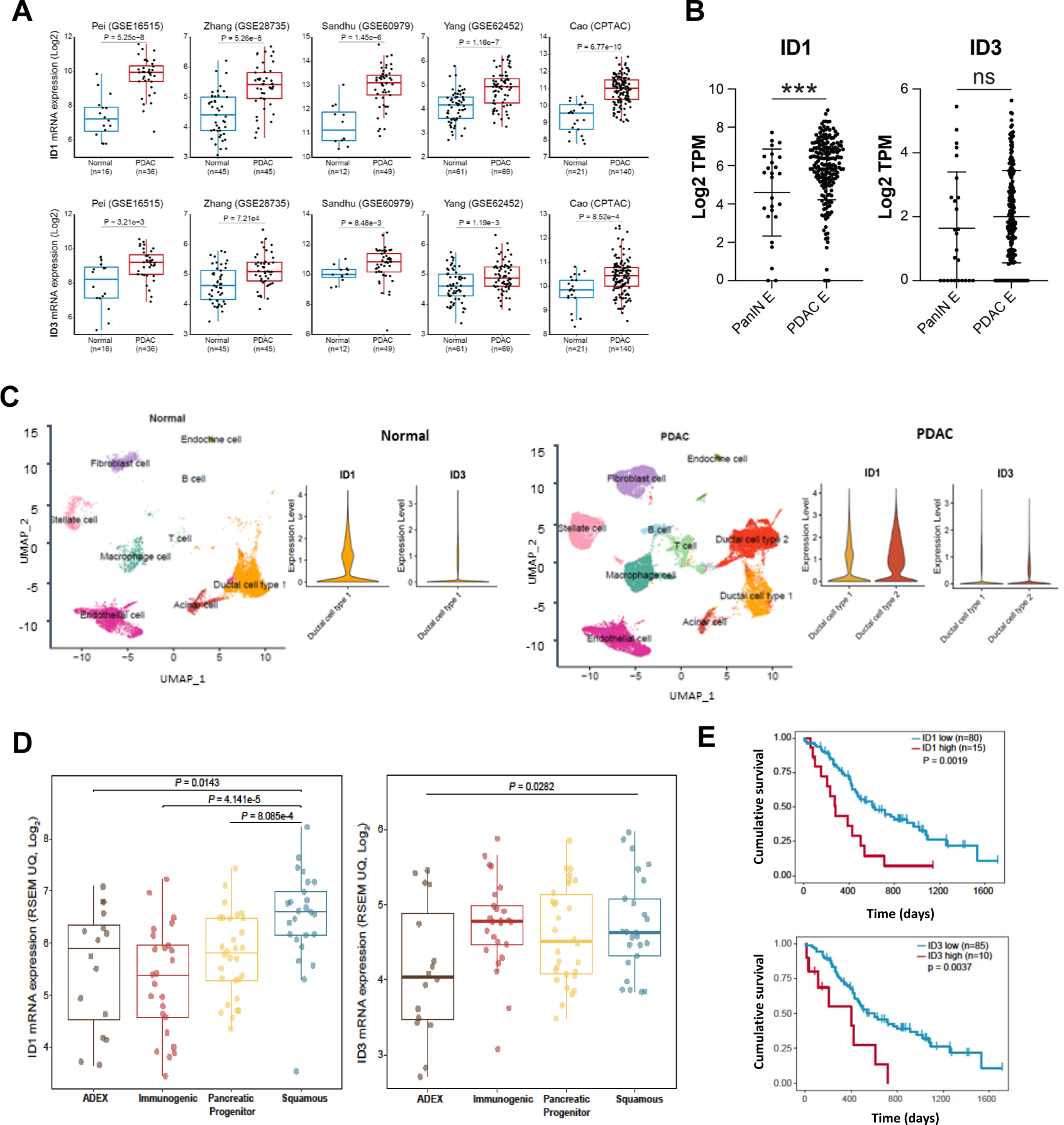
High expression of ID1 and ID3 in human PDAC is associated with shorter patient survival. **(A)** Box plots of ID1 and ID3 mRNA expression in five independent data sets profiling bulk RNA expression from normal human pancreas and PDAC tissues. GSE16515, GSE28735, GSE60979, GSE62452 are microarray analyses. CPTAC (NCI Clinical Proteomic Tumor Analysis Consortium) is RNA-seq analysis. *P*-values were determined by Wilcoxon Rank Sum test. **(B)** GSVA analysis of ID1 and ID3 mRNA expression from RNA-seq of epithelial cells obtained by laser capture microdissection (LCM) from 26 low grade human PanIN samples and 197 malignant primary human PDAC samples published by Maurer et. al. (PMID 30658994). The mean and +/-1 SD are shown. P-value determined by two-tailed *t-*test. \*\*\**p*<0.005. TPM is transcripts per million. **(C)** UMAP re-analysis of scRNA-seq data from 11 normal and 24 human PDAC samples published by Peng et. al. (PMID 31273297). Violin plots of ID1 and ID3 mRNA expression in the ductal cell type 1 population in normal samples (left) compared to the ductal cell type 1 and ductal cell type 2 populations in PDAC (right). Normalized expression levels were computed using Seurat. **(D)** Box plots of ID1 and ID3 mRNA expression in four PDAC subtypes in bulk RNA-seq data from 95 human tumors with high epithelial content (≥40%) published by Bailey et. al. (PMID 26909576). In the same work, these four subtypes were also identified in 232 pancreatic cancers using array-based mRNA expression profiles. ADEX is aberrantly differentiated endocrine exocrine. **(E)** Kaplan-Meier survival analysis of patients stratified by expression levels of ID1 and ID3 mRNAs in bulk RNA-seq data published by Bailey et. al. (PMID 26909576). Median survival was estimated using the Kaplan-Meier method and the difference was tested using the log-rank test. *P*-values <0.05 were considered statistically significant.

To look specifically at ID1 and ID3 expression in epithelial cells, we examined ID1 and ID3 mRNA levels in our existing RNA-seq dataset of laser capture, microdissected epithelial tissue from 26 PanIN (each low-grade) and 197 primary PDAC patient samples (**Figure 1B**)^18^. There is a statistically significant increase in expression of ID1 but not ID3 between PanIN and PDACs.

To identify specific epithelial cell sub-populations within tumors that expressed ID1 and ID3 mRNA, we interrogated an existing scRNA-seq dataset comprising 24 PDAC tumor samples and 11 control pancreata ^19^. The published analysis showed that acinar and ductal cell type 1 populations were present as expected in the normal samples. PDAC tumors also contained the ductal cell type 1 population, but computational analysis revealed that this population differed from that in normal tissue. In PDAC, the ductal cell type 1 population was enriched for genes associated with disease related terms (e.g. inflammatory response, cell adhesion and migration). Strikingly, unlike normal samples, PDAC tissue also contained a unique ductal cell type 2 population that was enriched for cancer-related functions (e.g. cell proliferation, migration, and hypoxia) (**Figure 1C**). In addition, the cancer-specific ductal cell type 2 population had higher expression of CEACAM1/5/6 and KRT19 which are markers associated with poor prognosis in PDAC ^19^.

Our independent re-analysis of the aforementioned scRNA-seq dataset identified the same nine clusters in normal samples - acinar, ductal cell type 1, endocrine, fibroblast, stellate, B cell, T cell, macrophage and endothelial populations - and the additional tenth ductal cell type 2 population in PDAC samples ^19^. ID1 transcripts, and to a lesser degree ID3 transcripts, were present in the ductal cell type 1 population in both normal and PDAC samples. The cancer-specific ductal cell type 2 population, however, showed a significant increase in ID1 transcripts as compared to the ductal cell type 1 population in either normal or PDAC samples (**Figure 1C**).

We next considered whether high expression of ID1 and ID3 was associated with specific PDAC subtypes defined by expression analysis that correlated with histopathological characteristics ^20^. Using unsupervised clustering of bulk RNA-seq data from 95 human PDAC tumors with high epithelial content (>40%) Bailey et al. identified four subtypes of tumors – aberrantly differentiated endocrine/exocrine (ADEX), immunogenic, pancreatic progenitor, and squamous ^20^. Using this dataset, we queried expression levels of ID1 and ID3 mRNA in each of the four PDAC subtypes. For ID1, significantly higher expression was observed in the squamous subtype than in the immunogenic, pancreatic progenitor, or ADEX subtypes. For ID3, similar expression was found in the immunogenic, pancreatic progenitor and squamous while lower expression was found in the ADEX subtype (**Figure 1D**).

Survival analysis of these same patients stratified by PDAC subtype showed median survival of 25.6 months with ADEX, 30 months with immunogenic, 23.7 months with pancreatic progenitor and only 13.3 months for patients with the squamous subtype of tumor ^20^. Separately, we carried out survival analysis stratified by high versus low expression of ID1 and ID3 mRNA. For both ID1 and ID3, high expression was correlated with shorter survival time from diagnosis. Patients with high ID1 expressing tumors had a median survival of 300 days versus those with low ID1 expressing tumors who had median survival of 650 days, while patients with high ID3 expressing tumors had a median survival of 400 days versus those with low ID3 expressing tumors who had median survival of 600 days (**Figure 1E**).

### BMP signaling is upregulated in human PDAC

Given that BMP signaling has been shown to upregulate ID1 and ID3 gene expression in multiple tissues and having established that high expression of ID1 and ID3 mRNA in PDAC tumors is associated with shorter patient survival times, we asked if BMP signaling played a role in upregulation of ID1 and ID3 in PDAC ^21–23^.

To investigate this, we examined the expression of BMP ligands and receptors in a large bulk RNA-seq data set (Pei et. al., GSE16515) and found that BMP2, BMP4 and the Type II receptor BMPR2 were all significantly upregulated in PDAC. In the same five datasets previously examined in Figure 1A ^12–16^, we observed that BMP2 was significantly upregulated in two while BMP4 and BMPR2 were significantly upregulated in PDAC tumor tissue in all five datasets (**Figure 2A**).

**Fig. 2:**
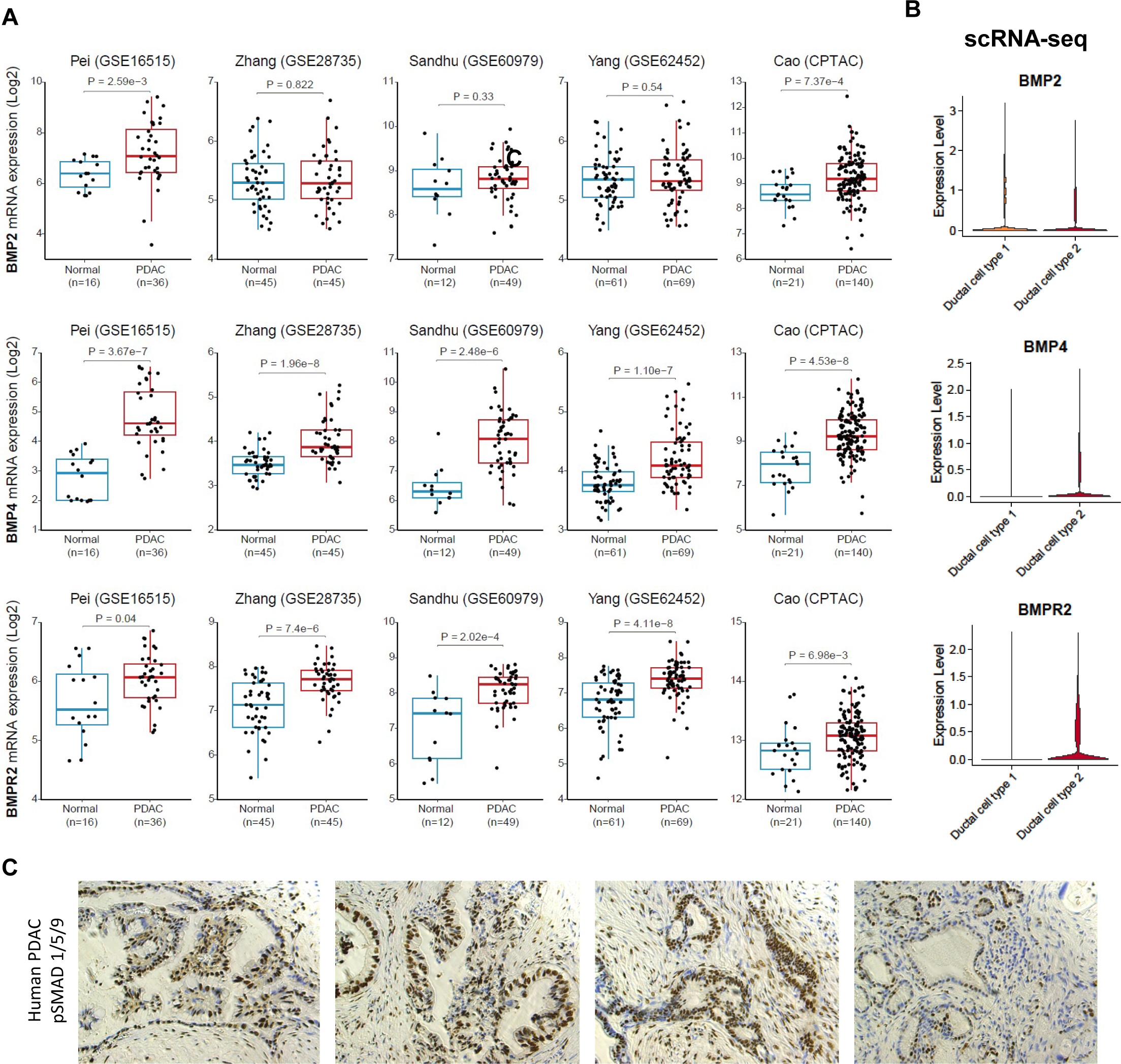
BMP signaling is upregulated in human PDAC. **(A)** Box plots of mRNA encoding BMP2 and BMP4 ligands and BMPR2 receptor in five independent data sets profiling bulk RNA expression from normal human pancreas and PDAC tissues GSE16515, GSE28735, GSE60979, GSE62452 are microarray analyses. CPTAC (NCI Clinical Proteomic Tumor Analysis Consortium) is RNA-seq analysis. *P*-values were determined by Wilcoxon Rank Sum test. **(B)** Violin plots of BMP2, BMP4 and BMPR2 mRNA expression in the ductal cell type 1 population and the ductal cell type 2 populations in PDAC using scRNA-seq data from 11 normal and 24 human PDAC samples published by Peng et. al. (PMID 31273297). Normalized expression levels were computed using Seurat. **(C)** pSMAD 1/5/9 immunostaining of FFPE pancreas tissue samples from four human PDAC patients.

Consistent with the finding that BMP4 and BMPR2 are the most highly and reproducibly upregulated BMP signaling molecules in PDAC tumors based on bulk RNA-seq analyses, we next analyzed their expression in a scRNA-seq dataset^19^. Here we observed upregulated expression of BMP4 and BMPR2 in the PDAC-specific pathogenic ductal cell type 2 population as compared to the ductal type 1 cell population which is also present in normal pancreas (**Figure 2B**) ^19^.

High levels of BMP4 and BMPR2 mRNA observed in the PDAC-specific ductal cell type 2 population led us to examine BMP-mediated SMAD signaling activity in PDAC lesions in human patient samples. A primary measure of active BMP signaling is nuclear localization of phosphorylated SMAD1/5/9 proteins (pSMAD1/5/9). Immunostaining of human PDAC tissue from four patients revealed high levels of nuclear pSMAD1/5/9 in the lesions of diseased pancreata, providing evidence of active BMP signaling in all four patient samples (**Figure 2C**).

### Upregulation of ID1 and ID3 mRNA expression by BMP signaling is blocked by Noggin and DMH1

To investigate a functional relationship between BMP signaling and expression of ID1 and ID3 in PDAC, two human PDAC cell lines were used: PANC1 and 779. PANC1 is an established epithelial carcinoma cell line while 779 is a patient-derived cell line isolated from a primary PDAC that we recently described ^24,25^. Both PANC1 and 779 cell lines harbor oncogenic mutations in *KRAS* and loss-of-function mutations in *TRP53*. They also lack expression of p16^INK4a^/CDKN2A but retain expression of SMAD4 ^25^.

As a first test of BMP signaling capacity, PANC1 and 779 cells were treated with 100 ng/mL BMP2 for 3 hours. Western blot analysis revealed increased levels of pSMAD1/5 protein and qRT-PCR analysis detected increased ID1 and ID3 mRNA in both PANC1 and 779 cells in response to BMP2 (**Figures 3A-C**). Conversely, treatment for 6 hours with 100 ng/mL Noggin, an endogenous regulator of BMP signaling which functions by binding to BMP ligands and preventing their interaction with BMP receptors, lowered pSMAD 1/5 protein in PANC1 and 779 cells relative to controls (**Figures 3A-B**). Noggin treatment also lowered ID1 mRNA in PANC1 cells, and ID3 mRNA in both PANC1 and 779 cells (**Figure 3C**). Similarly, treatment for 24 hours with 10 uM DMH1, a highly selective small molecule antagonist of the BMP receptors ALK2 (*ACVR1*) and ALK3 (*BMPR1A*), or with 3 nM LDN193189, a second small molecule antagonist of the same BMP receptors, decreased pSMAD 1/5 protein (**Figures 3A-B, S2**) ^26,27^. Importantly, DMH1 also significantly downregulated expression of ID1 and ID3 mRNA in both PANC1 and 779 cells (**Figure 3C**). Consistent with canonical BMP signaling occurring in PANC1 and 779 cells, Western blot analysis of cytoplasmic and nuclear fractions of BMP2-treated PANC1 cells revealed that higher levels of pSMAD1/5/9 induced by BMP2 was translocated to the nucleus (**Figure 3D**). Inhibition of BMP signaling by treatment with Noggin, DMH1 or LDN193189 demonstrates that BMP signaling is active in these cell lines and that blocking BMP signaling significantly lowers expression of ID1 and ID3 mRNA.

**Fig. 3:**
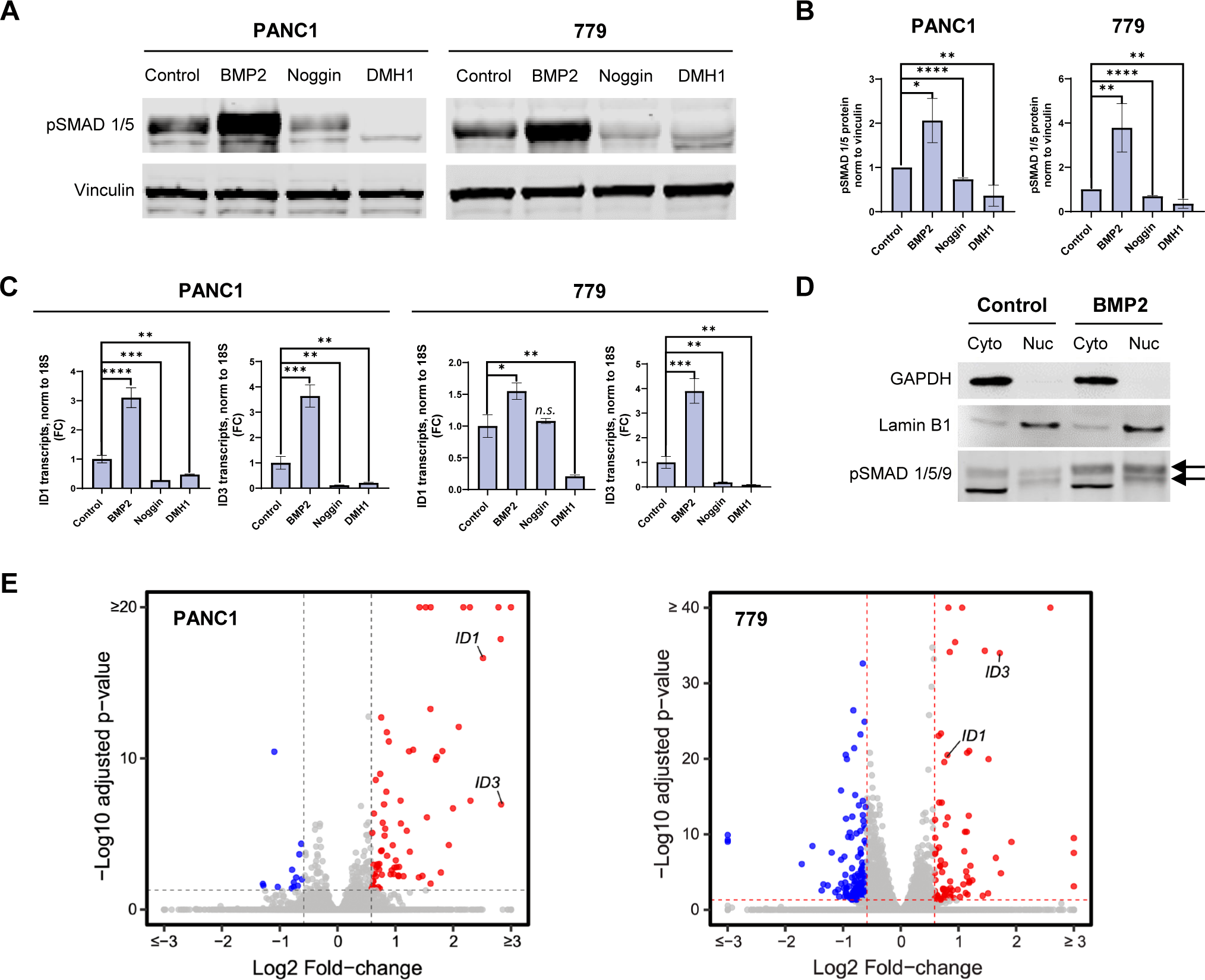
BMP signaling upregulation of ID1 and ID3 mRNA expression is blocked by Noggin and DMH1. **(A)** Western blot analysis of protein extracts from PANC1 and 779 cells following treatment with BMP2 (100 ng/mL), the BMP antagonist Noggin (100 ng/mL), or the highly selective small molecule ALK2/ALK3 BMP receptor inhibitor DMH1 (10uM). **(B)** Relative quantitation of pSMAD 1/5 in Western blots from three independent experiments in PANC1 and 779 cells treated with BMP2, Noggin or DMH1. In each experiment, signals were normalized to Vinculin. ImageJ (http://rsb.info.nih.gov/ij/) was used to calculate area and pixel value from images acquired using LI-COR Image Studio software. Error bars represent standard deviation. Statistical significance was determined by two-tailed *t* test. \**p* < 0.05, \*\**p* < 0.01, \*\*\*\**p* < 0.001. **(C)** RT-qPCR quantitation of ID1 and ID3 mRNA in human PANC1 and 779 cells following treatment with BMP2, Noggin or DMH1. Values were normalized to 18s rRNA. Error bars represent standard deviation. *P*-values were determined by two-tailed *t* test. \**p* < 0.05, \*\**p* < 0.01, \*\*\**p* < 0.005, \*\*\*\**p* < 0.001, *n.s.* not significant. **(D)** Western blot analysis of pSMAD1/5/9 in cytoplasmic versus nuclear fractions of protein extracts from human PANC1 cells following treatment with BMP2 ligand. GAPDH is a control for the cytoplasmic fraction and Lamin B1 is a control for the nuclear fraction. **(E)** Volcano plots with statistically significant log2 fold-changes in transcript levels in PANC1 and 779 cells following 3-hour treatment with BMP2 compared to control samples. Data is derived from two independent RNA-seq data sets for each sample in each cell line.

Having shown that BMP2 upregulates expression of ID1 and ID3 mRNA, we next sought to identify where ID1 and ID3 ranked in the hierarchy of BMP regulated target genes. RNA-seq analysis of PANC1 and 779 cells following 3-hour treatment with 100 ng/mL BMP2 showed that ID1 and ID3 mRNAs were among the most significantly upregulated target genes (**Figure 3E**). Together, the data suggest that a primary function of BMP signaling in PDAC is to increase expression of ID1 and ID3 which then disable control of gene expression by E2A or its splice variant E47.

### DMH1 downregulates genes that control the cell cycle

Based on the downregulation of pSMAD 1/5 protein and ID1 and ID3 mRNA observed in PANC1 and 779 cells following treatment with DMH1, we also conducted RNA-seq of both cell lines following 24-hour treatment with 10 uM DMH1. Consistent with the PCR data (**Figure 3C**), RNA-seq data from DMH1 treated cells showed downregulation of ID1 by 43% and ID3 by 75% in PANC1. In 779, DMH1 treatment reduced ID1 by 71% and ID3 by 90%. GSEA analysis of RNA-seq data revealed that two of the top three pathways identified in both cell lines were ‘G2M Checkpoint’ and ‘E2F Targets’. Pathway enrichment plots and heatmaps of the top 50 genes in each pathway are shown for both cell lines (**Figures 4A-D**). For the G2M Checkpoint pathway, 32 of the top 50 genes are the same in PANC1 and 779 cells (**Figure 4C**). For the E2F pathway, 28 of the top 50 genes are the same in PANC1 and 779 cells (**Figure 4D**). In agreement with the data identifying cell cycle control genes as the most highly downregulated targets by treatment with DMH1 in two PDAC cell lines, we also observed 20% fewer cells in DMH1-treated samples relative to control cultures after 24 hours (**Fig. 4E**). Taken together, the DMH1 studies reveal that active BMP signaling in PDAC plays a significant role in control of the cell cycle.

**Figure 4.**
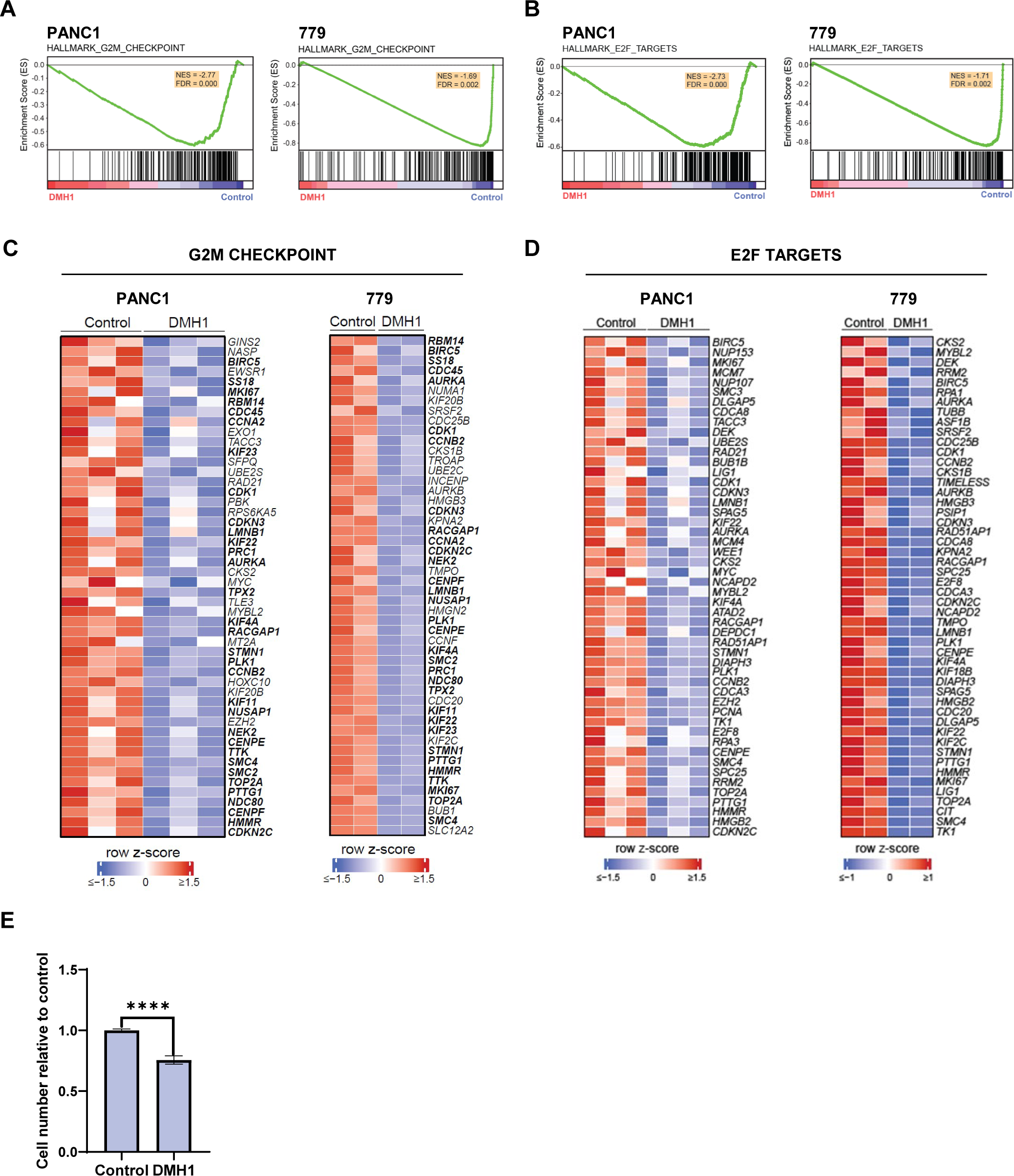
DMH1 downregulates genes that control the cell cycle. **(A)** GSEA enrichment plots of RNA-seq data from PANC1 and 779 cells following 24-hour treatment with DMH1 compared to controls using Hallmark G2M gene set. **(B)** GSEA enrichment plot of RNA-seq data from PANC1 and 779 cells following 24-hour treatment with DMH1 compared to controls using Hallmark E2F Targets gene set. **(C)** Heatmaps of the top 50 core-enriched genes following 24-hour treatment with DMH1 compared to controls according to GSEA analysis of the G2M gene set. **(D)** Heatmaps of the top 50 core-enriched genes following 24-hour treatment with DMH1 according to GSEA analysis of E2F Targets gene set. **(E)** Proportion of PANC1 viable cells following 24-hour treatment with DMH1 relative to control samples. Data is based on three independent experiments. Relative cell numbers were identified using alamarBlue staining measured on a fluorescence spectrophotometer using 560 nm excitation/590 nm emission filter settings. Statistical significance was determined by two-tailed *p-*test, \*\*\*\**p* < 0.001.

### BMP receptors, SMAD effectors, and ID1 and ID3 are upregulated in mouse models of early pancreas pathogenesis

Acinar cells in the pancreas can transdifferentiate to a progenitor-like ductal cell type through a process referred to as acinar-to-ductal metaplasia (ADM). ADM can be regenerative in cells that have experienced inflammation or injury but in cells expressing oncogenes such as *Kras^G12D^,* ADM progresses to pancreatic ductal neoplasia (PanIN) which can, in turn, progress to PDAC^28^. In six to eight week-old *Ptf1a^CreER^*, *LSL-Kras^G12D^*, *LSL-tdTomato* mice, injection of tamoxifen induced expression of oncogenic Kras^G12D^ in pancreatic acinar cells and labeled those same cells with tdTomato fluorescent protein. At six months post-tamoxifen injection, pancreata were collected and tdTomato+ cells were isolated by FACS for scRNA-seq. The reported analysis of the data on these lineage-labeled acinar cells showed that cells expressing oncogenic *Kras^G12D^* gave rise to metaplastic cells that expressed high levels of Id1 and Id3 mRNA compared to acinar cells^29^. Moreover, the authors reported that Id1 and Id3 were two of only five transcription factors that were highly expressed in metaplastic cells but showed very low or non-detectable expression in acinar cells.

To provide a window into an even earlier stage of pathogenesis marked by the appearance of ADM in the absence of oncogenic mutations, we injected adult *Ptf1a^CreERTM^;Rosa^LSL-EYFP/+^ (CERTY)* mice with tamoxifen to label pancreatic acinar cells with EYFP. After labeling, pancreatic injury was induced with repeated injection of the cholecystokinin ortholog, caerulein. Two to four weeks later, pancreata were collected and EYFP+ cells were isolated by fluorescence activated cell sorting (FACS) for scRNA-seq^30^. We then compared this single cell data from *CERTY* mice to scRNA-seq data from *Ptf1a^CreER^*, *LSL-Kras^G12D^*, *LSL-tdTomato* mice ^29^.

UMAP identified eight clusters in EYFP+ cells from mice with caerulein-induced injury and 11 clusters in tdTomato+ cells from Kras^G12D^ expressing mice. Three out of the 19 clusters – acinar, EEC and tuft - were common in samples from both types of mice. In Kras^G12D^ expressing mice, tdTomato+ acinar cells gave rise to tumor, metaplastic and senescent cells in the pancreas. In the injured pancreata, EYFP+ cells gave rise to ADM that comprised heterogeneous epithelial cells, primarily acinar-mucin-ductal and mucin-ductal sub-populations (**Figure 5A**).

**Fig. 5:**
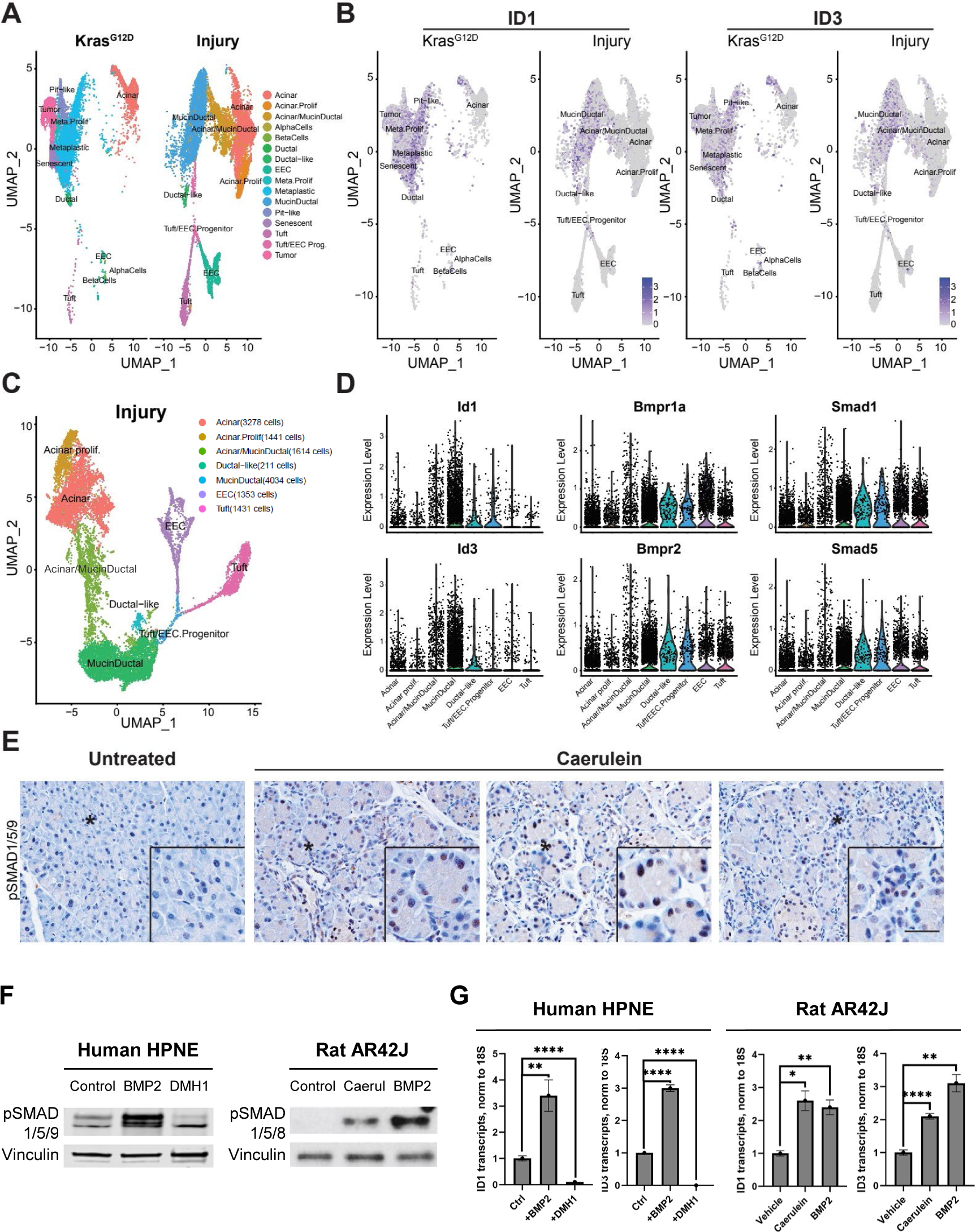
BMP receptors, Smad effectors, and Id1 and Id3 are upregulated in mouse models of early pancreas pathogenesis. **(A)** scRNA-seq UMAP annotated clusters of cells from adult mice harboring inducible *Ptf1a-Cre* to generate expression of *tdTomato* and *Kras^G12D^* (left) or *EYFP* (right) in pancreatic acinar cells. *EYFP*-expressing mice underwent subsequent caerulein-induced injury prior to isolation of EYFP+ cells for scRNA-seq analysis (right). **(B)** Cells expressing Id1 and Id3 mRNA are highlighted in purple within the UMAP annotated clusters shown in Figure 5A (gray). Key indicates the level of transcripts per cell. **(C)** Trajectory analysis showing annotated clusters in sorted cells from adult caerulein-induced injury mice harboring inducible *Ptf1a-Cre* to generate expression of *EYFP* (right) in pancreatic acinar cells. The order of the cell populations in the key, from top to bottom, follows the inferred developmental trajectory of the EYFP+ labeled acinar cells following caerulein-induced injury (Ma KDG Gastro 2022). **(D)** Violin plots showing mRNA expression of Id1, Id3, and BMP signaling molecules in the clusters shown in Figure 5C. Cluster types are shown from early to late (left to right) according to their inferred developmental trajectory (Ma KDG Gastro 2022). Normalized expression was computed using Seurat. **(E)** FFPE pancreas tissue from 4-week caerulein-treated and control mice immunostained for pSMAD1/5/9. **(F)** Western blot analysis of human HPNE cells following 3-hour BMP2 or 24-hour DMH1 treatment, and rat AR42J cells following 24-hour caerulein or 3-hour BMP2 treatment. **(G)** RT-qPCR quantitation of ID1 and ID3 mRNA in human HPNE cells following 3-hour BMP2 or 24-hour DMH1 treatment, and rat AR42J cells following 24-hour caerulein or 3-hour BMP2 treatment. Values were normalized to 18s rRNA. Error bars represent standard deviation. Statistical significance was determined by two-tailed *t* test. \*\**p* < 0.01, \*\*\*\**p* < 0.001.

We next interrogated expression of Id1 and Id3 mRNA in all populations of the Kras^G12D^ and injured mice pancreata. UMAP revealed that expression of Id1 and Id3 mRNA was highest in the tumor, metaplastic proliferating, metaplastic, senescent and pit-like cell populations of the Kras^G12D^ mice pancreata. In the injured pancreata, expression of Id1 and Id3 mRNA was highest in the mucin-ductal cells (**Figure 5B**). Together the data reveal that the appearance of all diseased cell populations arising from acinar cells, in response to oncogenic Kras^G12D^ or caerulein-induced injury, is coincident with induction of Id1 and Id3 mRNA expression.

Our previous trajectory analysis of the EYFP+ acinar cells from injured pancreata inferred their developmental progression from acinar cells through acinar-mucin-ductal to mucin-ductal cell identities (**Figure 5C**) ^30^. Strikingly, violin plots of Id1 and Id3 mRNA in each of these populations show that Id1 and Id3 expression rises with each progressive stage of pathogenesis. To further examine a possible link between Id1 and Id3 mRNA and BMP signaling in the earliest stages of pancreas pathogenesis, expression of *Bmpr1a*, *Bmpr2*, *Smad1* and *Smad5* was also quantified in each of the EYFP+ populations following injury. Compared to the acinar population, which serves as a normal control in this analysis, the mucin-ductal cell population contained the highest fraction of cells expressing BMP signaling molecules *Bmpr1a*, *Bmpr2*, *Smad1* and *Smad5* (**Figure 5D**). Consistent with this data, immunostaining of tissue from caerulein-treated mice revealed that the injured pancreata demonstrated a level of nuclear pSMAD1/5/9 expression not observed in untreated control tissue thereby providing evidence of active, canonical BMP signaling in the earliest pancreatic lesions (**Figure 5E**).

Having shown that caerulein treatment induced BMP signaling and expression of Id1 and Id3 mRNA in a murine model of pancreatitis, we wanted to determine whether this occurred in a cell-autonomous manner or required paracrine signaling from neighboring cells. We employed HPNE cells, a non-tumorigenic cell line derived from hTERT immortalization of human pancreatic exocrine cells, treated with BMP2 or DMH1. Data from Western blot analysis mirrored that of PDAC lines with increased pSMAD1/5/9 in response to BMP2 and decreased pSMAD1/5/9 in response to DMH1 (**Figure 5F**). RT-qPCR analysis showed that expression of ID1 and ID3 mRNA also increased in response to BMP2 and decreased in response to DMH1 (**Figure 5G**). Together, the data support the hypothesis that BMP signaling, response to inhibitors of BMP signaling, and expression of ID1 and ID3 mRNA can occur in a cell-type autonomous manner prior to tumorigenesis.

The rat acinar cell line AR42J has been widely used to study exocrine pancreas function particularly as an *in vitro* model of caerulein-induced acute pancreatitis ^31^. AR42J cells maintain many characteristics of normal pancreatic acinar cells including calcium signaling and synthesis and secretion of digestive enzymes. AR42J cells were treated with caerulein or BMP2, the later as a positive control. Western blot analysis showed that short term treatment with caerulein was sufficient to induce pSMAD1/5/8 (pSMAD8 is alternate nomenclature for pSMAD9) (**Figure 5F**). RT-qPCR analysis showed that expression of ID1 and ID3 mRNA also increased in response to both caerulein and BMP2 (**Figure 5G**). Together the data suggest a circuit in which exocrine cell stress, in this case modeled by treatment with caerulein, activates BMP signaling to induce expression of ID1 and ID3 in a cell-type autonomous manner.

## Discussion

The tumorigenic function of ID proteins is well established in many cancers ^32,33^. In PDAC, studies from our lab and others have clearly demonstrated that ID proteins play a pathogenic role and that knockdown of ID1 or ID3 decreases PDAC tumorigenicity ^6,7,9,10^. Our previous work showed that ID1 and ID3 bind to and inhibit the activity of the bHLH transcription factor E47, a splice variant of the *E2A*/*TCF3* gene. Importantly, we found that restoring E47 activity in PDAC induced quiescence and expression of acinar cell differentiation markers *in vitro* and *in vivo* ^7,25^. Given the increasing interest in IDs in PDAC, we investigated upstream signals that might control expression of the ID genes ^9,10,29^. BMP signaling was of particular interest as it’s known in many diverse biological processes to induce expression of ID1 and ID3 ^34^.

Here, we report that BMP signaling is active in human PDAC tumors as evidenced by the high expression of BMP ligands and receptors, and by induction of nuclear pSMAD1/5/9. BMP signaling upregulates expression of ID1 and ID3 which is, in turn, correlates with shorter patient survival times. We have determined that BMP signaling is required for high levels of ID1 and ID3 expression because blocking BMP signaling with either the endogenous antagonist, Noggin, or the small molecule BMP antagonists, DMH1 and LDN193189, reduced expression of both ID1 and ID3 mRNA relative to control samples. In agreement with the role of IDs in cell cycle control ^32^, 24 hours of treatment with DMH1 resulted in decreased cell number relative to controls and significant downregulation of genes encoding cell cycle related targets of E2F transcription factors and genes that control the G2M checkpoint. Conversely, treating cells with recombinant BMP2, which was previously shown to be highly expressed in PDAC, significantly increased expression of ID1 and ID3 indicating that expression in PDAC cell lines is sub-maximal under normal culture conditions ^35^. Further, our studies in human tissues and cells, as well as mouse tissue and rat cells, suggest that BMP signaling to IDs in the exocrine pancreas is conserved across species.

scRNA-seq analysis of PDAC patient samples established the emergence of two cancer cell populations in PDAC progression – one early epithelial ductal cell type 1 population and a second late-stage ductal cell type 2 population ^19^. Our independent analysis of human scRNA-seq datasets ^19^ from normal pancreas and PDAC also demonstrated significant upregulation of ID1 expression specifically in the ductal cell type 2 population. Subsequent trajectory analysis of the mouse PDAC GEMM datasets revealed that expression of Id1 was particularly elevated as malignancy progressed from the early-stage epithelial ductal cell type 1 population to the later-stage ductal cell type 2 population ^9^. The same authors analyzed a human scRNA-seq dataset comparing normal pancreas to PDAC and identified ID1 as the top upregulated transcription factor specifically in cancer cells and suggested a role for ID1 in EMT ^9^.

Although there has been litle work on the role of BMP signaling in pancreatic exocrine pathogenesis, the BMP inhibitor GREM1 was recently identified as a key regulator of cellular heterogeneity in pancreatic cancer. GREM 1 was highly expressed in the mesenchymal ‘EPCAM low’ tumor cell population where it acted as a paracrine factor to inhibit BMP signaling in neighboring epithelial ‘EPCAM high’ PDAC cells. Inactivation of Grem1 in a PDAC GEMM directly converted epithelial into mesenchymal PDAC while overexpression of Grem1 produced an epithelial phenotype in mesenchymal tumors ^36^.

With the significance of high expression of ID1 and ID3 in late-stage PDAC well-established, it was striking to observe that BMP signaling and induction ID1 and ID3 expression occur early in pancreas pathogenesis. In our experiments, caerulein treatment of mice induced development of diseased populations that transdifferentiated into increasingly abnormal ADM sub-populations with increasing expression of Id1, Id3 and effectors of BMP signaling. This finding is consistent with another study that also found high expression of pSMAD1/5/8 in the pancreata of caerulein-treated mice, although the authors did not evaluate expression of ID proteins. These authors went on to show that administration of Noggin blocked induction of pSMAD1/5/8 and reduced some features of caerulean-induced pancreatitis *in vivo* suggesting a therapeutic effect of inhibiting BMP signaling ^37^.

An unresolved question is how loss of SMAD4 (*DPC4* gene) affects BMP signaling to ID1 and ID3. SMAD4 is considered a required cofactor for canonical BMP signaling and TGFβ signaling. Inactivating mutations in SMAD4 are observed in approximately 55% pancreatic cancers, generally occurring in late lesions (PanIN-3) ^38^. Huang et. al. reported nuclear expression of ID1 in 87.5% of SMAD4 positive patient tumors and 71% of SMAD4 negative tumors (ID3 was not examined). Similarly, these authors observed ID1 staining in 46-47% of murine tumor cells from GEMM models regardless of SMAD4 status ^10^. Thus, regardless of SMAD4 mutational status, expression of ID1 is maintained throughout disease progression. In SMAD4 negative tumors, it will be of interest to determine whether BMP signaling continues to upregulate IDs through non-canonical pathways, or whether BMP independent pathways modulate IDs.

Pancreas cancer is projected to become the second leading cause of cancer-related mortality in the U.S. by the year 2030. With more than 90% of cases diagnosed after metastasis has occurred, current treatment regimens result in a relative 5-year survival of only 12.5% based on SEER data for 2013-2019. The interaction of ID proteins with bHLH transcription factors has been suggested as a therapeutic target in multiple cancers but the size of the protein-protein interface is relatively large and therefore notoriously difficult to target with small molecules. An alternative strategy may be to identify druggable signaling pathways that impinge on expression of ID1 and ID3 and/or their activity in cancer. BMP signaling is an atractive target for small molecule intervention as it is a highly druggable pathway.

In summary, our findings indicate that BMP signaling to genes that encode ID1 and ID3 is active at each stage of pathogenesis in the exocrine pancreas, providing insight into the etiology of pancreatic maladies.

## Materials and Methods

### Cell Culture

Human pancreatic cancer cell lines, PANC1 and 779 were cultured in RPMI-1640. 779 PDAC cell line was established from patient-derived tumor was by A.M. Lowy, M.D. of the Moores Cancer Center, UCSD. Human normal pancreatic cell line, HPNE was cultured in DMEM. All cell lines were purchased from ATCC and were cultured in media supplemented with 10% fetal bovine serum and 100 Units/mL Penicillin/Streptomycin antibiotics (Gibco, Catalog no. 15140-122). All cells were grown in 5% CO₂ at 37°C. 100 ng/mL BMP2 (Peprotech, Catalog no. 120-02C) was added to cells in culture media for 3 hours. 10 uM DMH1 (Fisher Scientific, Catalog no. 501364456) was added to cells in culture media for 24 hours. 100-200 ng/mL Noggin (Peprotech, Catalog no. 250-38) was added to cells in culture media for 6 hours.

### Quantitative RT-PCR

Total RNA was extracted from drug/vehicle-treated cell lines using NucleoSpin RNA kit (Takara, Catalog no. 740955.50). 2 ug of total RNA was used to synthesize cDNA with qScript Ultra Supermix (Quantabio, Catalog no. 95217). 1 uL of cDNA was used as template for qPCR. Quantitative RT-PCR used CFX96 Touch Real-Time PCR Detection System (BioRad) with PowerUp SYBR Green Master Mix (Applied Biosystems, Catalog No. A25741) with Quantitect human primers (Qiagen) for genes ID1, ID3 and 18S (used for normalization).

### Western blots

Treated cells were lysed in 2X Laemmli buffer (10% β-mercaptoethanol, 20% glycerol, 0.004% bromophenol blue, 0.125 M Tris HCl pH 6.8, 4% sodium dodecyl sulfate) heated for 10 minutes at 100°C and centrifuged at 17,000 x g for 15 minutes. Samples were analyzed by SDS-PAGE (4-12% Bis-Tris gel; Bio-Rad Laboratories, Inc.). Gels were transferred to nitrocellulose membranes, blocked in Intercept blocking buffer (Li-Cor, Catalog no. 927-70001), and incubated overnight at 4°C with primary antibodies listed below. Goat anti-rabbit (IRDye®-680RD RRID:AB_10974977) secondary antibodies were used at 1:5000 dilution for 1 hour at room temperature. After washes with 1xTBST, membranes were imaged on Licor Odyssey CLX with ImageStudio software. Band intensities were quantified and analyzed by ImageJ.

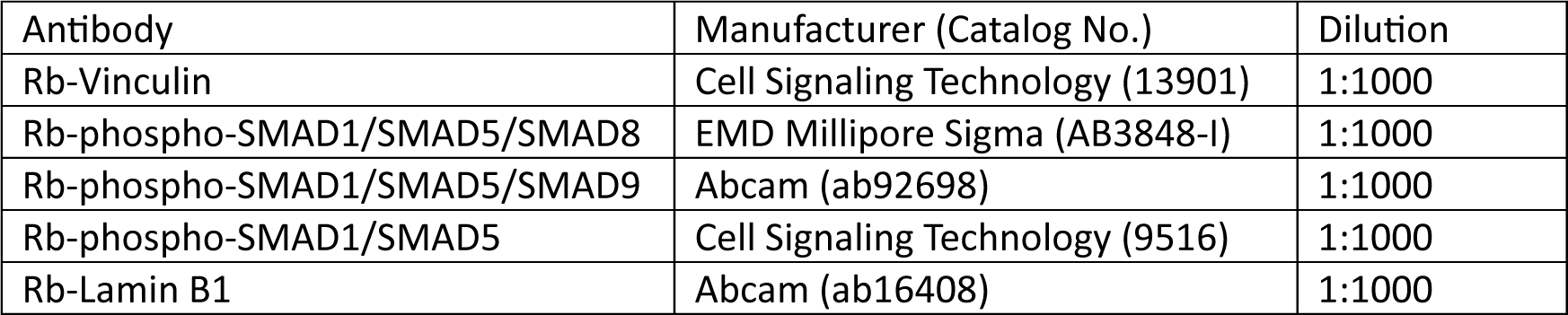

### Immunohistochemistry

Mouse pancreata were harvested and fixed in 4% PFA. Paraffin embedding, sectioning and slide preparations were done in the SBP Histopathology Core Facility. 5 uM sections on slides were stained on the automated IHC BONDRX (Leica) with anti-phospho-SMAD1/SMAD5/SMAD9 antibody (Abcam, ab92698) with antigen retrieval at pH 9.0. Goat anti-rabbit-HRP was used as the secondary antibody with the BOND polymer refine detection system (Leica, catalog no. DS9800). Images were taken by Zeiss LSM710 confocal microscope with a 10X, 20X and 40X objective lens.

### Gene expression comparison

Microarray datasets including normalized expression levels and metadata from GEO were downloaded in R v4.2.1 using Bioconductor package GEOquery v2.64.2. Gene expression levels were summarized from probe level to gene level by using the median of probes for a given gene. Probes mapped to multiple genes were discarded. Gene expression (RNA-seq) and clinical data from TCGA-PAAD Pan-Cancer 2018 study ^39^ were downloaded from cBioPortal ^40^. Datasets from Cao et. al. ^16^ were downloaded from NCI GDC Data Portal. Tumor versus normal sample comparisons were performed using Wilcoxon rank sum test in R.

### Analysis of published scRNA-seq datasets from Ma et. al

Processed count matrices for scRNA-seq datasets from Ma et. al. were downloaded from the Gene Expression Omnibus (GEO) database (accession number GSE172380). Low-quality cells were filtered on read counts, the number of genes expressed, and the ratio of mitochondrial reads following the thresholds described in the publication ^30^. Filtered gene count matrices were log-normalized, and the top 2000 variable features were further scaled prior to dimension reduction by PCA and being embedded in UMAP using the R package Seurat ^41^. Seurat cell clusters were labeled with major cell types using marker genes provided by the authors.

### Analysis of published scRNA-seq datasets from Peng et. al

Processed single cell count matrix and cell annotations were downloaded from Genome Sequence Archive under accession PRJCA001063. We used Seurat v4.0.5 for integration and analysis of the data ^42^. Raw count matrix and cell metadata were converted to Seurat object. Sample information (tumor or normal) was gleaned from cell barcodes. Raw data was normalized using the *NormalizeData* function. Top 2000 highly variable genes were selected and used for downstream analysis. Data was scaled using *ScaleData*. PCA on highly variable gene set was run using *RunPCA*. Data was integrated using Harmony v0.1.0 ^43^ based on sample information (tumor and normal samples). *RunUMAP*, *FindNeighbors*, and *FindClusters* functions were to used compute UMAP and cluster the cells using top 40 principal components and resolution of 0.5. Cell types were labelled as described by the original authors ^19^.

### RNA-seq

PolyA RNA was isolated using the NEBNext® Poly(A) mRNA Magnetic Isolation Module and barcoded libraries were made using the NEBNext® Ultra II™ Directional RNA Library Prep Kit for Illumina® (NEB, Ipswich MA). Libraries were pooled and single-end sequenced (1X75) on the Illumina NextSeq 500 using the High output V2 kit (Illumina Inc., San Diego CA).

### Bulk RNA-seq analysis

Raw reads were preprocessed by trimming Illumina Truseq adapters, polyA, and polyT sequences using cutadapt v2.3 (Martin, M., 2011). Trimmed reads were aligned to human genome version hg38 using STAR aligner v2.7.0d_0221 ^44^ and parameters according to ENCODE long RNA-seq pipeline (<u>htps://github.com/ENCODE-DCC/long-rna-seq-pipeline</u>). Gene expression levels were quantified using RSEM v1.3.1 ^45^. Ensembl v84 gene annotations were used for the alignment and quantification steps. RNA-seq sequence, alignment, and quantification qualities were assessed using FastQC v0.11.5 (htps://www.bioinformatics.babraham.ac.uk/projects/fastqc/) and MultiQC v1.8 ^46^. Batch effect correction was performed using *ComBat-seq* function in R package sva version 3.44.0 (Leek JT, Johnson WE, Parker HS, Fertig EJ, Jaffe AE, Zhang Y, Storey JD, Torres LC (2022) sva: Surrogate Variable Analysis, R package version 3.44.0). Lowly expressed genes were filtered out by retaining genes with estimated counts (from RSEM) ≥ number of samples multiplied by 5. Filtered and batch-corrected estimated read counts from RSEM were used for differential expression comparisons using the Wald test implemented in the R Bioconductor package DESeq2 v1.22.2 based on generalized linear model and negative binomial distribution ^47^. Genes with Benjamini-Hochberg corrected *p*-value < 0.05 and fold change ≥ 1.5 or ≤ -1.5 were selected as differentially expressed genes. RPKM values were computed from batch-corrected counts and gene lengths obtained from RSEM using edgeR package version 3.38.4 ^48^. Gene set enrichment analysis was done using GSEA app version 4.3.2. For PANC1 with three biological replicates for each of DMH1 and Control, we used *Run GSEA* with “Permutation type = gene set”. For 779 with two biological replicates for both DMH1 and Control, we ran pre-ranked GSEA using *Run GSEAPreranked.* Genes were ranked using adjusted p-values and log2 fold change by multiplying -Log10 of adjusted *p*-value and Log2 of fold change.

### Survival analysis

Survival analysis was performed in R version 4.2.1 using survival, survminer, and maxstat packages. Optimal cutpoints for the categorization of samples as ‘high’ and ‘low’ expressors of given gene in each dataset were determined using *surv_cutpoint* and *surv_categorize* functions from survminer package.

### Statistical analysis

Western blot and RT-qPCR data are indicated as Mean ± Standard Deviation. Statistical significance was evaluated by unpaired two-tailed Student’s t test. *P*-values are presented as *p<0.05, **p<0.01, ***p<0.001.

## SUPPLEMENTAL

**Figure S1.**
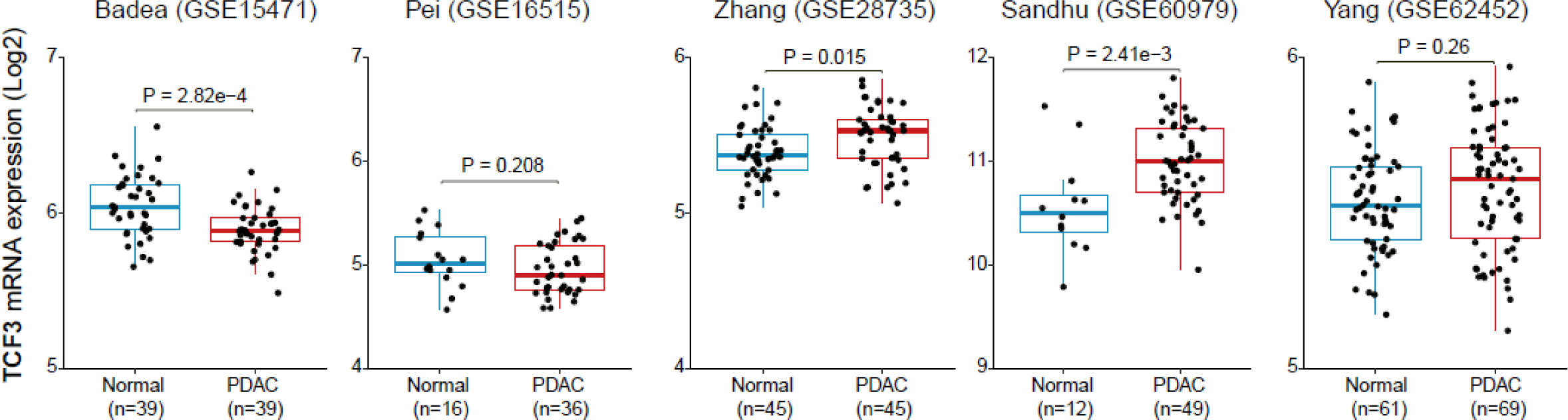
TCF3 (E2A) expression in human normal versus PDAC bulk RNA-seq samples. TCF3 (E2A) mRNA expression in five existing independent data sets profiling bulk RNA expression from human normal and PDAC tissues analyzed on microarrays. *P*-values were determined by Wilcoxon Rank Sum test.

**Figure S2.**
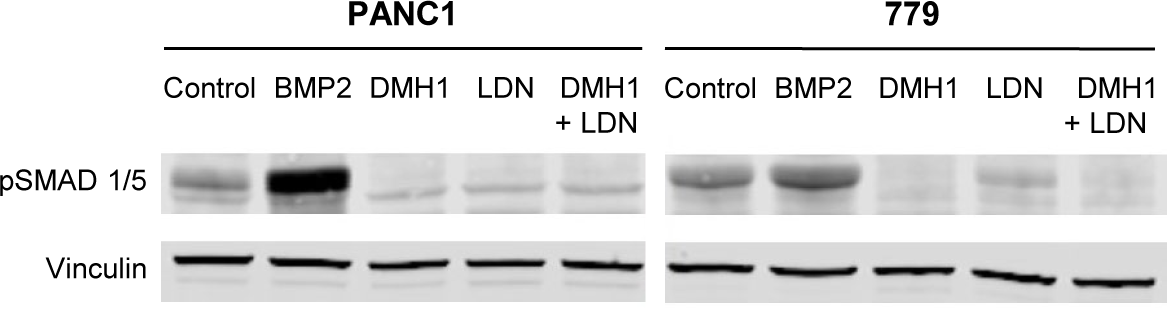
DMH1 and LDN193189 block phosphorylation of SMAD 1/5. Western blot analysis of pSMAD 1/5 protein in PANC1 and 779 cells following 3-hour treatment with BMP2 (100 ng/mL), or 24-hour treatment with the ALK2/ALK3 BMP receptor inhibitors DMH1 (10 uM), LDN193189 (3 nM), or DMH1 (10 uM) and LDN193189 (3 nM) combined.

**Figure S3.**
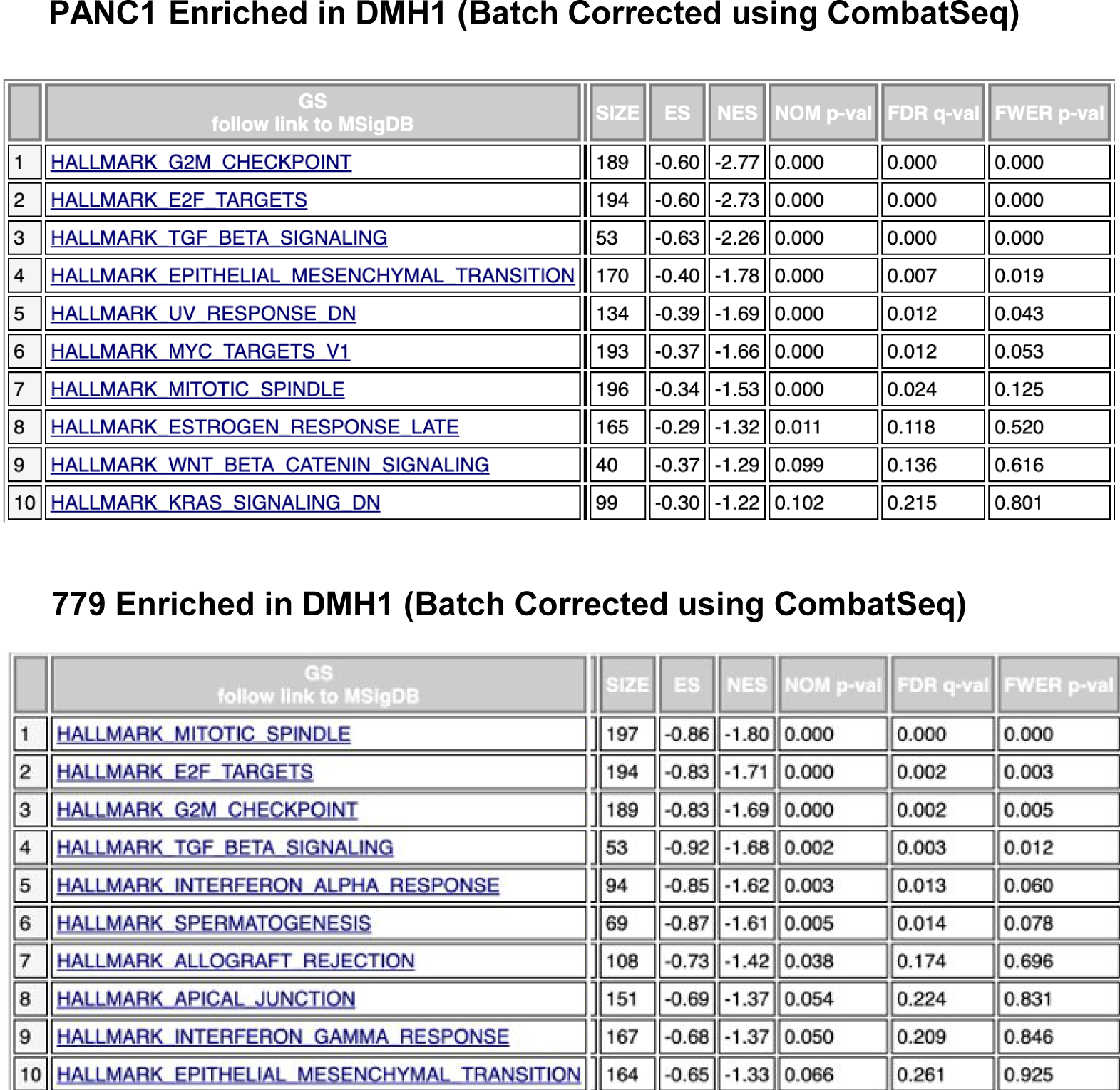
GSEA categories enriched in DMH1 treated samples relative to control samples in PANC1 and 779 cells. The results of DEseq analysis following batch correction using CombatSeq. Using *Run GSEAPreranked,* genes were ranked using adjusted *p*-values and Log2 fold change by multiplying -Log10 of adjusted *p*-value and Log2 of fold change.

## Notes

### Competing Interest Statement

The authors have declared no competing interest.

### Summary of Updates

Two authors added - Rabi Murad and H. Carlo Maurer. Additional data added.

